# M_t_ or not M_t_: Temporal variation in detection probability in spatial capture-recapture and occupancy models

**DOI:** 10.1101/2023.08.08.552394

**Authors:** Sollmann Rahel

## Abstract

State variables such as abundance and occurrence of species are central to many questions in ecology and conservation, but our ability to detect and enumerate species is imperfect and often varies across space and time. Accounting for imperfect and variable detection is important for obtaining unbiased estimates of state variables. Here, I investigate whether closed spatial capture-recapture (SCR) and single season occupancy models are robust to ignoring temporal variation in detection probability. Ignoring temporal variation allows collapsing detection data across repeated sampling occasions, speeding up computations, which can be important when analyzing large datasets with complex models. I simulated data under different scenarios of temporal and spatio-temporal variation in detection, analyzed data with the data-generating model and an alternative model ignoring temporal variation in detection, and compared estimates between these two models with respect to relative bias, coefficient of variation (CV) and relative root mean squared error (RMSE). SCR model estimates of abundance, the density-covariate coefficient β and the movement-related scale parameter of the detection function σ were robust to ignoring temporal variation in detection, with relative bias, CV and RMSE of the two models generally being within 4% of each other. An SCR case study for brown tree snakes showed identical estimates of density and σ under models accounting for or ignoring temporal variation in detection. Occupancy model estimates of the occupancy-covariate coefficient β and average occupancy were also largely robust to ignoring temporal variation in detection, and differences in occupancy predictions were mostly <<0.1. But there was a slight tendency for bias in β under the alternative model to increase when detection varied more strongly over time. Thus, when temporal variation in detection is extreme, it may be necessary to model that variation to avoid bias in parameter estimates in occupancy models. An occupancy case study for ten bird species with a more complex model structure showed considerable differences in occupancy parameter estimates under models accounting for or ignoring temporal variation in detection; but estimates and predictions from the latter were always within 95% confidence intervals of the former. There are cases where we cannot or may not want to ignore temporal variation in detection: a behavioral response to detection and certain SCR observation models do not allow collapsing data across sampling occasions; and temporal variation in detection may be informative of species phenology/behavior or for future study planning. But this study shows that it can be safely ignored under a range of conditions when analyzing SCR or occupancy data.

## Introduction

State variables such as abundance and occurrence of species are central to many questions in ecology and conservation. It is well understood that our ability to detect individuals (e.g., Otis et al., 1978; Pollock et al., 1990; Williams et al., 2002) or species (e.g., MacKenzie et al., 2002; Tyre et al., 2003) in field surveys is imperfect (i.e., we may undercount individuals and fail to detect species even if they are present). Further, detection often varies across survey locations, methods, and conditions. Ensuring that important sources of variation in detection are accounted for is important for robust inference about ecological parameters (Pollock et al., 2002; Gimenez et al., 2008) and the development of methods that can estimate abundance while accounting for imperfect detection has a long history (Lincoln, 1930). The past decades have seen the development of a variety of hierarchical models (HM) that formulate sub-models for the (imperfectly observed) state and the conditional (on the state) detection process (Royle & Dorazio, 2008; Kéry & Royle, 2015). Essentially a combination of multiple Generalized Linear (Mixed) Models (GL(M)Ms), describing variation in detection is straight forward in HM.

Closed capture-recapture, both spatial (e.g., Royle et al., 2014) and traditional (e.g., Otis et al., 1978; Pollock et al., 1990), and occupancy modeling (MacKenzie et al., 2017) are popular frameworks to estimate animal abundance and occurrence, respectively, that explicitly model the state and detection process and thus allow for accounting for imperfect and varying detection. Closed population capture-recapture models use individual-level encounter data collected over multiple sampling occasions to estimate abundance while correcting for imperfect detection probability of individuals. The simplest estimator is model M_0_, which assumes that detection, *p*, is the same for all individuals at all times, which is unlikely in reality. There are three main sources of variation in *p* in closed CMR models (Otis et al. 1978): *p* can vary by individual, depend on whether an individual has been caught before, or across sampling occasions. The respective estimators accounting for these sources of variation are referred to as model M_h_ (heterogeneity), M_b_ (behavior) and M_t_ (time) (more than one source of variation in *p* can be present, and estimators can account for multiple such sources). When individual heterogeneity is present but not accounted for, estimates of abundance will be biased low as the sample will be skewed towards more detectable individuals. When ignoring behavioral responses to capture, bias in abundance can be positive or negative, depending on whether animals are trap shy or trap happy. Temporal variability in *p* can be induced by, for example, sampling-related factors such as sampling effort (e.g., Agresti, 1994), environmental conditions interacting with catchability of the study species (e.g., Wegan et al., 2012), or random variation. But contrary to behavioral and individual variation, Otis et al. (1978) showed via simulations that M_0_ was largely unbiased when data violated the assumption of constant detection over time.

Compared to traditional closed CMR, spatial capture recapture (SCR) explicitly describes the spatial nature of sampling and populations (Efford, 2004; Borchers & Efford, 2008; Royle & Young, 2008), thereby accounting for individual movement on and off the sampling grid and variation in exposure to sampling among individuals. In SCR, detection probability is modeled on the detector level as a declining function of distance to an individual’s activity center; thus, in addition to individual, behavioral and temporal variation, detection can vary in space (for example, due to setup location, Sollmann et al., 2011; or due to different trap types, Kervellec et al., 2023). Investigation of SCR estimator bias related to detection has largely focused on spatial variation (Moqanaki et al., 2021), as well as mis-specification of the detection function (Dey et al., 2022). In developing SCR models with sampling effort adjustments, Efford et al. (2013) showed that ignoring temporally varying effort that is consistent across detectors (i.e., induces only temporal but no spatial variation in detection probability) had negligible effects on density estimates. This suggests that SCR should be robust to violation of the assumption of time-constant detection. But to my knowledge, this has not been investigated more generally.

Whereas CMR models deal with individual detection probability when estimating abundance, single season occupancy models estimate species presence/absence accounting for species detection probability. Models are fit to repeated species detection/non-detection data collected across multiple sampling locations; thus, detection probability can be modeled as varying in space, as well as in time. Similar to SCR models, temporal (or spatio-temporal) variation in *p* can be the consequence of varying sampling effort (e.g., Kéry & Royle, 2009), environmental conditions affecting sampling efficiency (e.g., noise, Wiest & Shriver, 2016), but also of fluctuations in abundance over time (Royle & Nichols, 2003), for example due to migratory behavior that can make a species essentially non-detectable in parts of the survey (e.g., Flanders et al., 2015). Occupancy models have been compared, conceptually, to non-spatial capture recapture models in which the all-0 encounter histories (here, sampling locations without any detections) are observed (Nichols & Karanth, 2002). In spite of that similarity, I am not aware of any studies exploring the sensitivity of occupancy model estimates to ignoring temporal variation in *p*.

Here, I explore via simulation and case studies whether estimates of parameters of the ecological (or state) sub-model of SCR and occupancy models are robust to violation of the assumption of time-constant detection. Fitting SCR models in which detection is allowed to vary over time requires calculating the likelihood of each individual by trap by occasion data point (i.e., using 3D data). In contrast, when detection is constant over time, data for common observation models (Poisson, Binomial) can be summarized across occasions in a 2D format. A 3D data format can greatly increase computation time, which can become cumbersome or problematic in complex models and/or when working with large data sets. As for SCR, when *p* varies over time in occupancy models, site and occasion specific data must be used, whereas if it is constant over time, data can be summarized to binomial counts. Although generally far less time consuming than SCR models, complex occupancy models (i.e., with multiple random effects, for communities, combining multiple surveys, etc.), can take hours to days to fit and require large amounts of memory. As large scale and long term camera trap and other survey data sets increasingly become available, so will the demand for fitting complex models to large data, across study areas or time periods. If temporal variability in detection can be ignored safely, this will greatly contribute to keeping these models manageable and viable, including for users without access to designated high power computing resources.

## Methods

### Simulations

Both SCR and occupancy models (OM) estimate a partially latent state – density and occurrence – while accounting for imperfect detection of individuals (SCR) or species (OM), *p*. I implemented a simulation study to test the effect of ignoring temporal variation in detection on estimates of parameters associated with the ecological state (for model-specific details, see following sections; for an overview over all model-specific scenarios, see Table 1).

**Table 1.** Overview of simulation scenarios to investigate effects of ignoring temporal variation in detection probability *p* for occupancy models (OM) and spatial capture-recapture (SCR) models. ‘Specific scenario’ columns show which variations (if any) of the main scenarios were fit for each modeling framework. Input parameters for all scenarios can be found in Appendix 1.

| Main scenario | General description | Specific scenarios SCR | Specific scenarios OM* |
| --- | --- | --- | --- |
| 1 | Temporal variation in $p$ | 1 | 1 high<br>1 low |
| 2 | Spatio-temporal variation in $p$ | 2<br>2a: amount of temporal variability in $p$ varies in space<br>2a-10: same as 2a but with stronger temporal variability in $p$<br>2b: temporal variability in $p$ is skewed<br>2c: temporal variability in $p$ is autocorrelated | 2 high<br>2 low |
| 3 | Spatio-temporal variation in $p$, correlated with spatial variation in density/occupancy | 3 | 3 high<br>3 low |

\* high and low refer to average 6-fold and 3-fold variation in $p$ over time, respectively

For both frameworks, I explored three main scenarios of temporal variation in detection, by making it a logit-linear function of a covariate, *Z*. In scenario 1, *Z* increased linearly over time (sampling occasions) but was constant for all detectors. Here, *Z* could represent day/month of sampling in a study where all detectors operate at the same time; or the time since application of lure/bait if it was applied to all detectors once at the beginning of a study.

In scenario 2, the detector-level mean of *Z*, 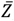, varied randomly across detectors following a standard normal distribution, and for each detector, occasion specific values of *Z* varied around 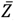 following a normal distribution with standard deviation SD=0.5. Possible examples for such spatio-temporally varying covariates might be activity of other species affecting space use of the focal species; varying sampling effort (e.g., in volunteer-run surveys where effort cannot be fully standardized, or due to malfunctioning or staggered setup/take-down of detector equipment; but note that in many cases, including effort as a form of offset may be preferred, Efford et al., 2013); or varying survey conditions, especially when not all locations can be surveyed simultaneously due to logistic constraints.

In scenario 3, I also generated a spatio-temporally varying *Z*, but one that was correlated with a spatial covariate affecting density/occupancy *X* (see below). Scenario 3 represents a situation in which not accounting for spatial variation in detection is likely to lead to bias in estimates of the state-covariate relationship.

For SCR models, I further created several variations on scenario 2: In scenario 2a, I randomly assigned detectors to one of 4 standard deviations for *Z* to mimic different degrees of variability in detection over time. In scenario 2b, I randomly assigned detectors to one of five Normal distributions with different amounts and directions of skew to create a situation in which the mean of *Z* is not necessarily the best representation of the most common conditions. Finally, in 2c, I created temporally correlated values of *Z* for each detector, as spatial correlation in detection probability, when ignored, has been shown to lead to bias in parameter estimates (Moqanaki et al., 2021).

Details of how *Z* was generated in all scenarios and for both frameworks can be found in Appendix 1. In both frameworks, I refer to models that account for temporal variation in detection only as M_t_, models that account for spatial variation only as M_s_, models that account for variation in both space and time as M_st_, and models that do not account for either as M_0_. In contrasting models that do vs do not account for temporal variation (i.e., M_t_ vs M_0_; M_st_ vs M_s_), I will refer to the latter as the alternative model.

#### SCR

For all scenarios, I simulated activity centers of *N*=60 individuals in a discrete state space with 24x26 cells, each measuring 50 by 50 units. I generated a spatially correlated covariate *X* for all 624 grid cells, and set the effect of *X* on density, *β* = 1. I randomly assigned activity centers to state space cells using a multinomial distribution with cell probabilities ***π* = *β*X/ Σ *β*X**. I generated detections across a 7x8 detector grid (*J*=56 detectors) spaced 100 units and centered in the state space, across *K*=8 sampling occasions. With this configuration, the state space extended 300 units beyond the trap array. I calculated detection probability of each individual *i* at each detector *j* and occasion *k* following a half-normal detection function,

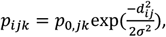

where *d*_*ij*_ is the distance between the activity center of individual *i*, **s**_*i*_, and the detector *j*, and

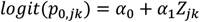

That is, in the SCR framework spatio-temporal variability in detection is modeled as variation in the baseline detection probability *p*0. Note that *Z*_1*k*_ = *Z*_2*k*_ = … = *Z*_*jk*_ in scenario 1. I set the scale parameter σ = 80 to ensure that individuals would be detectable at multiple detectors and chose values of α_0_ and α_1_ that led to similar overall detectability for all scenarios (with, on average, 20 individuals being detected at least once in all scenarios; for details on α_0_ and α_1_, see Appendix 1). I then generated individual, location and occasion specific detections as a Bernoulli random variable with *p*_*ijk*_.

With my settings for α_0_ and α_1_, variability in *p*_0_ at a given detector over time was approximately 7-fold for scenario 1, 3-fold for scenarios 2 and 3, and 6-fold in scenarios 2a, b and c. To check that the magnitude of variation in *p*_0_ was not driving results, I repeated scenario 2a (which showed the largest amount of bias in parameter estimates under the alternative model, see Results) with, on average, 10-fold variation in *p*_0_.

For each scenario, I generated 100 data sets and analyzed each data set with the data generating model (i.e., properly accounting for temporal variability in *p*_0_; M_t_ in scenario 1 and M_st_ in scenarios 2 and 3) and the corresponding alternative model ignoring temporal variation in *p*_0_ (M_0_ in scenario 1, M_s_ in scenarios 2 and 3). To fit model M_s_, I used the detector-level average value of *Z*, 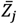, as a covariate on *p*_0_. All models used the half-normal detection function and included the effect of *X* on density.

I implemented models in the R package secr ver 4.5.6 (Efford, 2022), in R ver. 4.2.1 (R Core Team, 2022). For each model I kept track of model convergence, as well as estimates of all model parameters, and AICc; and I calculated realized abundance across the state space using the region.N() function. I considered a model as not converged either when likelihood maximization failed or when at least one standard error could not be calculated due to a singular Hessian matrix.

I assessed the effect of ignoring temporal variation in *p*_0_ by comparing median relative bias 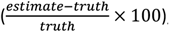, precision (coefficient of variation 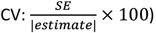), and accuracy (relative root mean squared error) between the data generating model and the corresponding alternative model for *N, σ* and *β*. These are typically the parameters of main ecological interest; *σ* is a detection parameter but is linked to animal movement, and as a detection parameter, could be more sensitive to mis-specification of the detection model. I opted for the median across datasets instead of the mean to minimize the effect of outlier iterations with extreme estimates. If SCR models are robust to violation of the assumption of constant *p*_0_ over time, I expect bias to be similar whether or not the model accounts for temporal variation in *p*_0_. Similar precision between the data generating and the alternative model would indicate similar coverage of truth. Finally, I calculated how often AICc suggested the data-generating model as the most parsimonious model, to gauge whether my simulations created sufficient temporal variation in *p*_0_ to be detectable in the data. I only counted a model as the most parsimonious when all other models had a ΔAICc≥2.

#### Occupancy

I implemented an analogous simulation study for occupancy models. I generated detection/non-detection data across *J*=100 detectors and *K*=8 sampling occasions. I modeled occupancy probability *ψ*_*j*_ as a function of a single, spatially correlated covariate *X*,

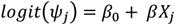

with *β* = 1 and *β*_0_ = 0.5. This corresponded to an average occupancy probability 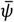 = 0.62. Temporal variation in detection probability was modeled as

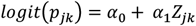

In all scenarios, α_0_ and α_1_ were chosen so that the total detector-level detection probability across all 8 sampling occasions was, on average, approximately 0.67 (for details, see Appendix 1). I explored only the main scenarios 1-3, as results from scenarios 2a-c from the SCR simulation suggested that these were not fundamentally different from main scenario 2 (see Figure 1, Appendix 2: Table S2). However, exploratory occupancy simulations suggested that sensitivity of occupancy models to ignoring temporal variation in *p* may depend on the magnitude of variability in *p* over time. I therefore explored two versions for all main scenarios, one where per-detector variability in *p* was, on average, 3-fold (labelled ‘low’), and one where it was 6-fold (labeled ‘high’).

**Figure 1.**
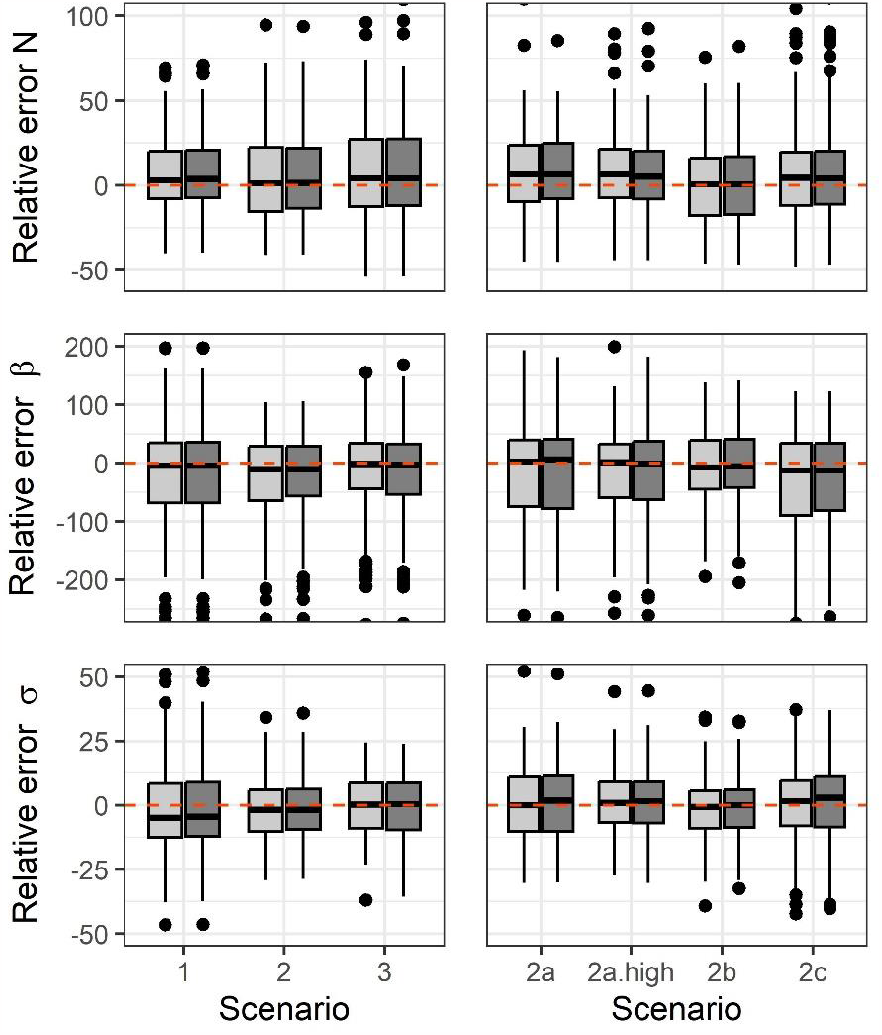
Relative error (boxplot) and bias (median) in estimates of abundance (N), covariate effect on density (β) and scale parameter of the half-normal detection function (σ) from data-generating spatial capture-recapture (SCR) model accounting for temporal variation in baseline detection probability *p*_0_ (light-grey) and alternative SCR model not accounting for temporal variation in *p*_0_ (dark-grey), for 100 simulated data sets. Red dashed line shows zero error/bias. In scenario 1, *p*_0_ varies over time, in 2 and 3, over time and space. Scenarios 2 and 2a-c differ in degree and shape of temporal variation in *p*_0_. In scenario 3, detection and density covary. For scenario details, see main text and Appendix 1.

As occupancy models were much faster to fit, I ran 250 iterations per scenario, using the package unmarked ver. 1.2.5 (Fiske & Chandler, 2011). I used the same criteria for convergence as described above. A few converged models had parameter estimates >|10|, with similarly large or much larger standard errors. Because in a real analysis with scaled covariates such estimates would be considered unreasonable, I considered iterations with any parameter estimate >|10| as not converged. For all converged iterations, I compared relative bias, CV and RMSE for estimates of *β* and 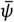 (i.e., *logit*^-1^(*β*_0_), as *X* was centered on 0) between the data-generating and the alternative model. I further calculated the maximum difference in occupancy predictions across sampling locations, as well as the number of sampling locations with an absolute difference >0.1. As for SCR, I tallied how often the data-generating and alternative model were selected as the most parsimonious model by AIC (the default criterion in the package unmarked).

### Case studies

#### SCR – brown tree snakes

I re-analyzed the brown tree snake data set described in Amburgey et al. (2021a) collected along 27 transects in a fenced area inside Andersen Air Force Base, Guam. Data are freely available from Amburgey et al. (2021b). All transects were split into 16-m segments, resulting in 351 potential detection locations. In November and December 2016, researchers conducted 32 night time surveys during which transects were either sprayed with scent the day of the survey (fresh) or the previous day (old), to determine whether scent improved detectability of snakes. As application of the two scent treatments varied in space and time, this is an example of scenario 2, with spatio-temporal variation in *p*_0_. In the original analysis, an SCR model was fit estimating *p*_0_ separately for no, old, and fresh scent, but results indicated that *p*_0_ was similar for no and old scent. I therefore grouped these two treatments into a single category, thus creating a binary predictor (no/old scent vs fresh scent). I fit model M_st_ using the location and occasion specific scent treatment, and model M_s_, for which I calculated the proportion of active sampling occasions with fresh scent for each location. I fit both models in secr, compared them by AICc, and compared estimates of density and *σ* between the two models.

#### Occupancy – Swiss breeding birds

I analyzed data for ten bird species (Appendix 3: Table S1) from the 2014 Swiss Breeding Bird survey (Schmid et al., 2004), available with the R package AHMbook v. 0.2.6 (Kéry et al., 2022). Data were collected 2-3 times along fixed transects during the breeding season by skilled volunteer birdwatchers across 266 1-km squares. I reduced the original count data to binary detection/non-detection data. The dataset provides several square and survey-specific covariates; following Kery and Royle’s (2015) community occupancy analysis of the full dataset, I considered elevation (m), elevation squared and percent forest cover as predictors of occupancy, and Julian date (1 April = 1), date squared and survey duration (min) as predictors of detection (M_st_). For the model accounting only for spatial variation in *p* (M_s_), I also calculated average survey date and duration for each square. I scaled all covariates prior to analysis. As average date was highly correlated with elevation (*r*=0.84), this is an example of scenario 3. The 2014 dataset consists of 145 species with detections. I first selected species with >50 detections (n=77) to avoid sparse data issues, then fit the above described model M_st_ to data from each of these 77 species. I predicted detection probabilities across all squares and occasions based on that model and calculated the ratio between the maximum and minimum *p* at each square. I kept the ten species for which that ratio was, on average, >2, to avoid species with very little temporal variation in *p*. For these species, I also fit model M_s_, with the same structure on occupancy, and average date, average date squared and average duration on *p*. I compared both models with respect to AIC, occupancy coefficient estimates **β** (relative difference (*β*_*Ms*_ - *β*_*Mst*_)/*β*_*Mst*_; and *β*_*Ms*_ within 95%CI of *β*_*Mst*_), model predictions of occupancy for sampled locations, 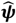 (maximum of 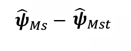 and 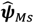 within 95%CI of 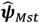), and response curves for individual occupancy covariates (curve under M_s_ within 95%CI for curve under M_st_).

## Results

### Simulations

#### SCR

Both the data generating model and the model ignoring temporal variation in *p*_0_ converged for all iterations in all scenarios. Even for these small simulated data sets, model run time increased almost 7-fold (from approximately 26 seconds to 179 seconds) from model M_s_ to model M_st_.

In all scenarios and for all three parameters, median relative bias (Figure 1), precision (CV; Appendix 2: Figure S1) and relative RMSE (Appendix 2: Table S2) were near-identical between the data-generating and the alternative model ignoring temporal variation in *p*_0_. Median relative bias under the two models was always within 4% of each other, and mostly within 1 or 2%, with the highest difference in bias being exhibited by *β* in both versions of scenario 2a (amount of variation in *p*_0_ over time varies across detectors). Differences in median CV and relative RMSE were similarly in the 1-3% range. The only exception was the RMSE for estimates of *β* in scenario 3, which was 8% higher under the alternative than the data-generating model.

According to AICc, the data-generating model was the most parsimonious model in 67% (scenario 2) to 93% (scenario 2c; 97% for 2a 10-fold) of all iterations. The corresponding model ignoring temporal variation was the most parsimonious model in 1-4% of all iterations (Appendix 2: Table A1). Thus, temporal variation in *p*_0_ translated to considerable temporal variation in detections in the data in almost all cases.

#### Occupancy

Both the data generating model and the model ignoring temporal variation in *p* converged in 230 - 249 iterations across all scenarios (Appendix 2: Table S3). Model run time was about 3 times higher for models accounting for temporal variation in *p*.

When detector-level temporal variation in *p* was, on average, 3-fold, for both parameters median relative bias (Figure 2), precision (CV; Appendix 2: Figure S2) and relative RMSE (Appendix 2: Table S4) were near-identical between the data-generating and the alternative model ignoring temporal variation in *p*, differing by at most 2%. The only exception was the estimate of *β*, which in scenario 1 had a 5% higher relative bias, and in scenario 2 had a 4% higher RMSE under the alternative model. When detector-level temporal variation in *p* was, on average, 6-fold, median relative bias, CV and relative RMSE in estimates of both parameters remained very similar for both models (generally within 4% of each other). But there was a tendency for *β* to show higher relative bias under the alternative model for all scenarios (7% higher for scenario 1, 5% higher for 2 and 3).

**Figure 2.**
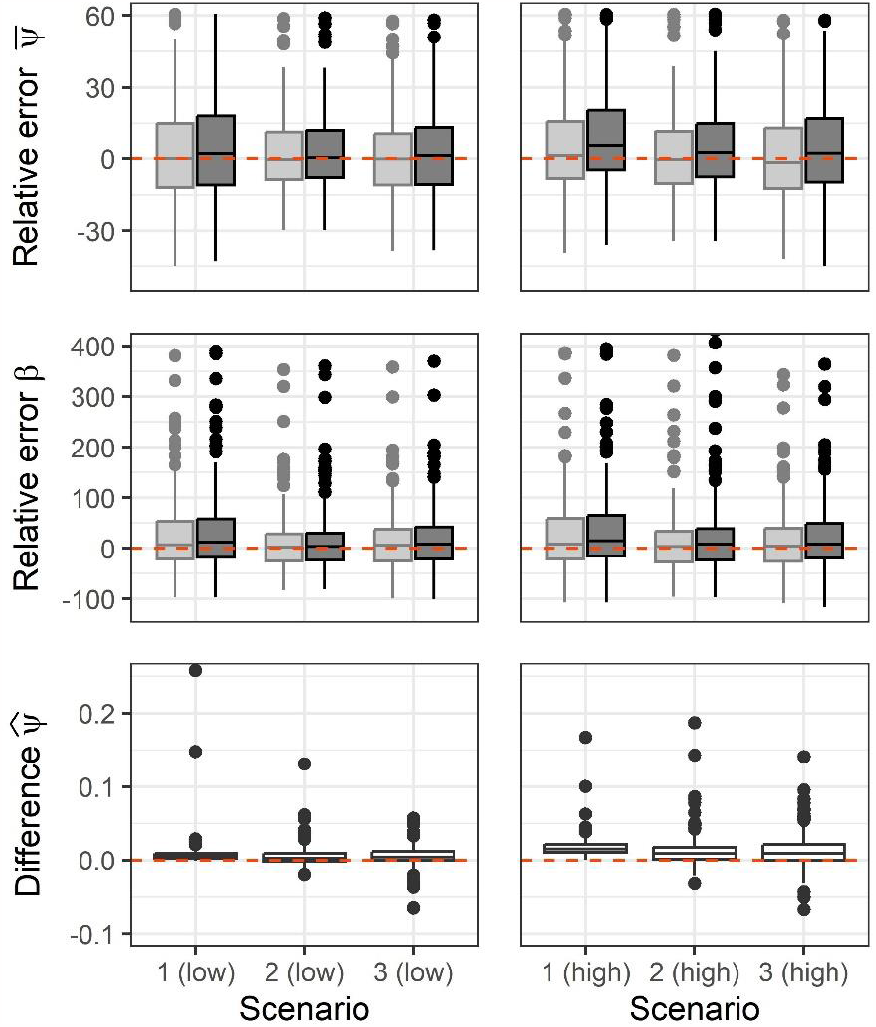
Relative error (boxplot) and bias (median) in estimates of mean occupancy 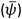 and covariate effect on occupancy (*β*) from data-generating occupancy model accounting for temporal variation in detection probability *p*_0_ (light-grey) and alternative model not accounting for temporal variation in *p*_0_ (dark-grey), and maximum difference in occupancy predictions 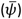 between these models, for 100 simulated data sets. Red dashed line shows zero error/difference. In scenario 1, *p*_0_ varies over time, in 2 and 3, over time and space. In scenario 3, occupancy and detection covary. Average variation in *p* over time either approximately 3-fold (low) or 6-fold (high). For scenario details, see main text and Appendix 1.

Corresponding to these minor differences in parameter estimates, the maximum difference in occupancy predictions for sampling locations between the data generating and alternative model was, on average across simulation iterations, 0.02 or lower for all 3-fold and 6-fold scenarios (Figure 2). The average number of sampling locations (out of 100) with an absolute difference >0.1 was 2.26 (scenario 3, 6-fold variation) or lower (Appendix 2: Table S4).

According to AIC, the data-generating model was the most parsimonious model in 83% (scenario 1, 6-fold) to 100% (scenario 1, 3-fold) of all iterations. The corresponding model ignoring temporal variation was the most parsimonious model in 1-3% of all iterations (Appendix 2: Table A3).

### Case studies

#### SCR – brown tree snakes

For the SCR analysis of the brown tree snake data, model M_st_ was clearly favored by AICc over model M_s_ (ΔAICc = 6.3). Estimates of *p*_0_ under M_st_ were 1.4×10^−3^ with no/old scent and 0.9×10^−3^ with fresh scent. Nonetheless, estimates of density and *σ* were indistinguishable (*D*: 22.24 individuals/ha ± 2.33, *σ*: 41.21 m ± 2.18 under both models). Model M_st_ took about half an hour to run. When compressing data into binomial counts for model M_s_, run time decreased to about 10 minutes.

#### Occupancy – Swiss breeding birds

For the occupancy analysis of ten Swiss breeding birds, M_st_ was clearly favored over M_s_ by AIC for 8 species (ΔAIC for M_s_ >2), comparable to M_s_ (ΔAIC for M_s_ <2) and less supported than M_s_ (ΔAIC for M_st_ >2) for one species each. Average site-level ratios of maximum to minimum *p* ranged from 2 to 89 (in the latter case, the species was essentially non-detectable early and late in the season; Appendix 3: Table S2, Figure S1).

Even though for several species, the relative difference in occupancy coefficients was considerable (>10% and up to 192% in one case of a coefficient close to 0; Appendix 3: Table S3), all coefficients under M_s_ were always within the 95%CI of the respective coefficients under M_st_. For all species, occupancy predictions at sampled locations under M_s_ were slightly higher than predictions under M_st_, with maximum differences per species ranging from 0.02 to 0.14. All predictions under M_s_ were always within the 95%CI of the respective predictions under M_st_ (Appendix 3: Figure S2). Similarly, all response curves for individual predictors under M_s_ were within the 95%CI envelope for the respective curves under M_st_, but curves reflected the tendency of M_s_ to lead to higher occupancy estimates (Figure 3). Parameter estimates for all ten species under both models can be found in Appendix 3: Table S3 and S4.

**Figure 3.**
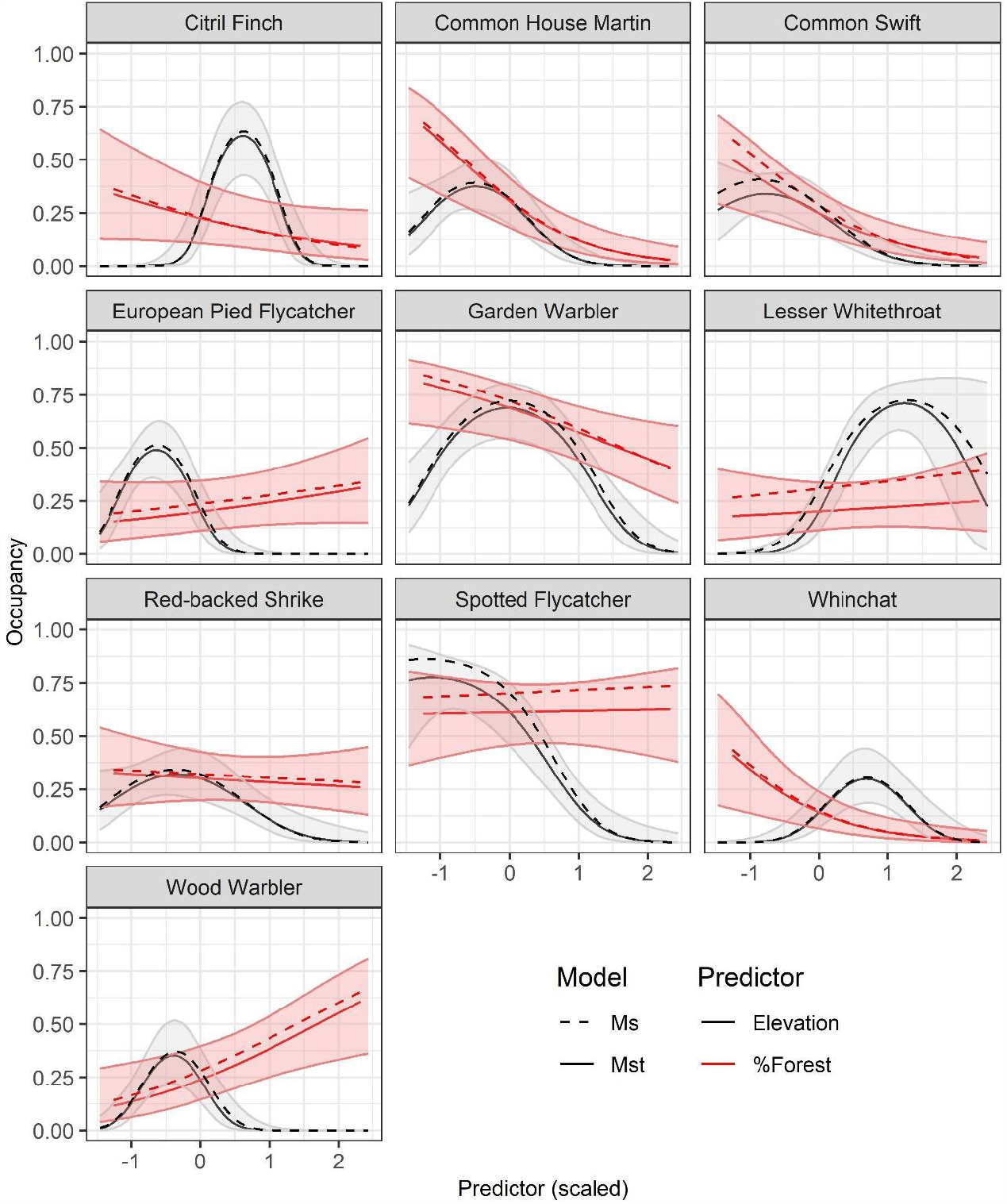
Relationship of occupancy with elevation and forest estimated with an occupancy model that does (M_st_) and one that does not (M_s_) account for temporal variation in detection, for 10 species of birds surveyed across Switzerland in 2014. Shaded areas are 95% confidence intervals of predictions under M_st_.

## Discussion

Based on simulation and case studies, I found that estimates of parameters of ecological interest from SCR and occupancy models were largely robust to ignoring temporal variation in detection probability. Compared to SCR, occupancy models appeared to be somewhat more sensitive to this assumption violation, particularly when temporal variability in *p* was high. However, even when estimates differed between occupancy models accounting for and ignoring this temporal variation, inference under both models about occupancy-covariate relationships and occupancy probabilities remained largely the same.

## Spatial capture-recapture

Similar to traditional closed CMR models (Otis et al., 1978; Williams et al., 2002), ignoring temporal variation in *p*_0_ in SCR models had little effect on estimates of abundance, the density-covariate relationship, or space use (σ). As such, this study provides yet more evidence that density estimates from SCR models are robust to various forms of mis-specification of the detection model. They have been shown to be robust to unmodeled spatial heterogeneity in detection probability, so long as that heterogeneity is not spatially autocorrelated (Efford et al., 2013; Bischof et al., 2017; Moqanaki et al., 2021), and to a range of mis-specifications of the detection function (Efford, 2004; Royle et al., 2014; Dey et al., 2022). In the latter case, though, inference about σ can be strongly affected by choosing the wrong detection function (Dey et al., 2022). In contrast, I found that estimates of σ were robust to ignoring temporal variation in detection. As σ in most applications relates to a stationary activity center and is modeled as constant across occasions, it appears sufficient to take into account spatial variation in *p*_0_ to obtain unbiased estimates of σ. It is conceivable that in closed SCR models where activity centers are allowed to move (Royle et al., 2016), not accounting for temporal variation in *p*_0_ may affect estimates of σ. Exploring this was beyond the scope of this study.

Beyond density, I found that estimates of the density-covariate relationship were not affected when temporal variation in *p*_0_ was ignored, so long as spatial variation was properly modeled. Efford et al. (2013) showed that ignoring spatial variation in sampling effort (and hence, detection) can bias estimates of this relationship. Though the effect of ignoring spatial variation in *p*_0_ was not the focus of the present simulation study, I did fit model M_0_ to data generated under scenarios 2 and 3 (i.e., with spatio-temporal variation in detection in data generation; results not shown). Abundance and σ estimates showed as little bias as under models M_s_ and M_st_, as did the density-covariate relationship in scenario 2 (no correlation between detection and density). But in scenario 3, where density and detection covaried, estimates of the density-covariate relationship under M_0_ had a median relative bias of -95%; as expected, in such situations variation in detection and density is confounded.

Even though different SCR simulation scenarios corresponded to different amounts of temporal variation in *p*_0_, results were very similar across scenarios, including for scenario 2a, which I ran with two levels of temporal variation in *p*_0_. This indicates (a) that SCR is robust to ignoring temporal variation in *p*_0_ across a range of magnitudes of such variation; and (b) that the lack of effect of violating the constant-*p*_0_ assumption is unlikely to be the result of being too conservative (with respect to temporal variation in *p*_0_) when generating data. The latter is supported by AICc identifying, in most cases, temporal variation in detection as important to explain variation in the data. Checking the raw generated data, even in scenarios with 3-fold variation in *p*_0_, the number of detections declined from about 7 in high-*p*_0_ occasions to about 2 in low-*p*_0_ occasions. In the brown tree snake case study, *p*_0_ was 50% higher on occasions when no fresh scent was present, and ignoring that variation had no effect on estimates of density or σ. Royle et al. (2014), based on another case study in which ignoring temporal variation in *p*_0_ had no effect on estimates of other parameters, suggested that the observed 3-fold variation was too weak to cause bias when ignored. But based on the present study, I suggest that temporal variation does not matter for obtaining unbiased estimates of these parameters, at least in the conditions explored here (but see the final paragraph for situations where ignoring temporal variation in detection may not be an option).

## Occupancy

Even though generally low, there was a tendency for bias in estimates of the occupancy-covariate relationship to increase in the simulation study when temporal variability in detection was high and not accounted for in the model. Moreover, when ignoring temporal variation in *p*, predicted occupancy probability was slightly higher than when accounting for it (that was the case in both the simulation and the case study). Maximum differences in simulations were typically <<0.1 and whether such small differences in site level occupancy predictions are relevant will depend on the study objective. Compared to simulations, differences in estimates and predictions between the two models were larger in the bird case study (though never statistically different based on 95%CI), but there was no clear pattern that these were related to the degree of temporal variation in detection. For example, of the five species with the largest amount of temporal variation in *p* (an average maximum-to-minimum ratio of *p* across occasions >10), three had a maximum difference in occupancy predictions >0.1, but the other two were comparable to the remaining species (maximum-to-minimum ratio of *p* <8), which had maximum differences ≤0.08. Ten species is too small a sample size to draw conclusions on the relationship between temporal variation in *p* and its effects on estimates when it is ignored. Still, I suggest that when extreme variability in *p* is suspected (e.g., due to extreme variation in survey effort or conditions, or due to known migratory or other behavior affecting species detection) and highly unbiased predictions of occupancy are important for the study objective, modeling temporal variability in *p* may be necessary. In contrast, if the general nature of relationships between occupancy and predictors is the focus, and/or extreme temporal variability in *p* is not expected, results suggest that temporal variability in *p* can be safely ignored.

In the bird case study, relationships of detection with predictor variables were sometimes markedly different, depending on whether that variable was included at the occasion level (for M_st_) or summarized across occasions (for M_s_). Conceptually, this is not surprising, as the two versions of the detection covariates measure different things – variation in space and time vs variation in space only. This may have implications, however, when AIC is used for variable selection, which is a common procedure in wildlife research (e.g., Murtaugh, 2009; Arnold, 2010). Thus, while fitting model M_s_ may not lead to appreciably different ecological inference when using the same covariate structure as one would use under M_st_ (i.e., when the detection model is determined a priori), inference about which covariates are important drivers of detection and occupancy may change if model selection is performed with data that are collapsed across occasions. Exploring the implications of modeling binomial counts instead of binary detections for model selection was beyond the scope of this study.

Finally, I suspect that the larger discrepancies in estimates/predictions between the two models in the bird case study are related to the more complex real world situation, including potential (albeit weak) correlation among multiple predictors in the model, as well as missing predictors and/or random variation not reflected in the model. It is possible that a similarly complex SCR data set with multiple predictors varying in space and time may also show larger discrepancies in parameter estimates between a model with and without temporal variation in detection.

## M_t_ or not M_t_: considerations for both modeling frameworks

The data and models underlying both simulation studies were simple enough (out of necessity) that fitting models M_t_ or M_st_ was trivial in terms of computation time; and even for the SCR case study, when fit in secr, it only took 30 minutes to run a model using the 3D data. But researchers increasingly leverage multiple and/or large datasets to answer questions on large temporal and spatial scales. In such circumstances, the more complex nature of the data and question often requires a custom model, often written in the BUGS language and implemented in a Bayesian framework. For example, Bischof et al. (2020) used seven years of data collected across much of Sweden and Norway to estimate spatial population dynamics of four carnivores, each with >6000 detected individuals, with SCR models. Tremendous computational advances have made it possible to fit SCR models to such large data at all. Vectorization, for example, – performing the same set of model computations on a vector, which is possible in nimble (de Valpine et al., 2017) – reduces memory usage and run time of SCR models (Turek et al., 2021). Vectorizing model computations over the (vector of) occasion-specific detection histories can counteract some of the added computational expense of 3D data. Nimble provides several other avenues to increase efficiency and speed up model fitting (Turek et al., 2021). Still, even with these improvements, models such as the one fit to the multi-year Swedish/Norwegian carnivore data remain extremely computationally challenging (Bischof et al., 2020). Having to use 3D data with projects of this scope can be a real limitation for what can be achieved in a reasonable time frame (noting that, while computation time in theory should not be a reason for not fitting the most appropriate model, in practice, projects and people operate under real time constraints).

When computation time is not a limiting factor, there is, of course no loss in fitting a model accounting for temporal variation in detection. When there is only temporal and no spatial variation in detection, then, the model ignoring temporal variation (M_0_) has one fewer parameter than the model accounting for that variation (M_t_). There was no indication in simulation results from scenario 1 that this reduction in model complexity led to a change in parameter precision. It is possible that in sparse data situations, the simpler model M_0_ may improve precision of parameter estimates, or may even be the only model that can be fit. On the other hand, for situations where spatio-temporal predictors of detection are summarized across occasions, we effectively lose information on the detection-covariate relationship and precision of estimates may suffer. Again, I found no indication of that in the simulation study, but for the bird case study, there was a slight tendency for standard errors of estimates to be higher under M_s_ than under M_st_ (Appendix 3: Table S4). Whether that matters will again depend on the specific data and objectives of a study.

There are also situations in which we have little choice but to fit a model taking into account the temporal dimension of the data. The first is by necessity: Some SCR observation models cannot be reduced to two-dimensional data, for example, the multinomial model in which an animal can only be detected in a single trap in a given occasions (Royle et al., 2014). Also in SCR models we may have to account for a behavioral response to being captured (or otherwise detected). Such a behavioral response effectively creates individual heterogeneity in detection over time, and ignoring such heterogeneity will bias estimates of density (e.g., Royle et al., 2014). A similar situation can occur in occupancy models, when detection of the species improves the observer’s ability to detect it again on subsequent visits (e.g., Riddle et al., 2010). The second situation is by design, when we are explicitly interested in temporal variation in detection. In SCR models, when data on other species is available at detectors, as is the case for camera traps, these can be used to study the spatio-temporal response in site use of the focal species to sympatric species (e.g., Bahaa-el-din et al., 2016). The same is true for occupancy models that take into account the detections of potentially interacting species when modeling detection of a focal species (e.g., Richmond et al., 2010). Further, in occupancy, temporal changes in detectability can be used to study species phenology, e.g., when birds are detected by song, which is indicative of territorial behavior and mate attraction during the breeding season (Strebel et al., 2014; Furnas & McGrann, 2018). Temporal variation in detection can also inform the design of futures studies, by revealing the most suitable times or conditions (i.e., associated with high detection probability) for sampling (e.g., Chambert et al., 2012; Amburgey et al., 2021a). As a side note, this study only refers to variation in detection across a set of secondary sampling occasions all nested within the same primary sampling period (in the simulation and case studies, only a single primary period is considered). In studies with multiple primary sampling occasions, variation in detection among primary occasions should be modeled, as otherwise, changes in detectability over time will be confounded with changes in abundance or occurrence (e.g., Pollock et al., 2002). Still, in many cases where detection is a mere nuisance variable and behavioral responses to capture/detection are unlikely (as is often the case with non-invasive survey methods), this study suggests that temporal variation in *p* across secondary occasions can be ignored under a range of conditions without meaningful impact on estimates of density or occupancy and their correlates.

## Appendices

Appendix 1: Input parameters for simulation scenarios

Appendix 2: Detailed results of simulation study

Appendix 3: Detailed results of occupancy case study of Swiss breeding birds

Appendices can be found at https://doi.org/10.5281/zenodo.8221229 (Sollmann, 2023a).

## Acknowledgements

I would like to thank the members of the Team Biodiversity Dynamics in the Department of Ecological Dynamics (IZW) for their feedback on this study. I also thank the EURING 2023 audience, and specifically Cyril Milleret for constructive questions that helped refine this study. Finally, I thank Ben Bolker, Ben Augustine and an anonymous reviewer for their constructive feedback on this manuscript.

## Data, scripts, code, and supplementary information availability

Scripts and code can be found at https://doi.org/10.5281/zenodo.10053390 (Sollmann, 2023b). The article only uses previously published data.

## References

Agresti A (1994) Simple capture-recapture models permitting unequal catchability and variable sampling effort. Biometrics, 494–500. 10.2307/2533391

Amburgey SM, Yackel Adams AA, Gardner B, Lardner B, Knox AJ, Converse SJ (2021a) Tools for increasing visual encounter probabilities for invasive species removal: a case study of brown treesnakes. NeoBiota, 70, 107–122. 10.3897/neobiota.70.71379

Amburgey SM, Lardner B, Knox AJ, Converse SJ, Yackel Adams AA (2021b) Brown Treesnake detections on transects using potential attractants of live-mouse lures or fish-spray scent, Guam: U.S. Geological Survey data release. 10.5066/P9G6JHZ3

Arnold TW (2010) Uninformative Parameters and Model Selection Using Akaike’s Information Criterion. The Journal of Wildlife Management, 74, 1175–1178. 10.1111/j.1937-2817.2010.tb01236.x

Bahaa-el-din L, Sollmann R, Hunter LTB, Slotow R, Macdonald DW, Henschel P (2016) Effects of human land-use on Africa’s only forest-dependent felid: The African golden cat Caracal aurata. Biological Conservation, 199, 1–9. 10.1016/j.biocon.2016.04.013

Bischof R, Milleret C, Dupont P, Chipperfield J, Tourani M, Ordiz A, de Valpine P, Turek D, Royle JA, Gimenez O (2020) Estimating and forecasting spatial population dynamics of apex predators using transnational genetic monitoring. Proceedings of the National Academy of Sciences, 117, 30531–30538. 10.1073/pnas.2011383117

Bischof R, Steyaert SM, Kindberg J (2017) Caught in the mesh: Roads and their network-scale impediment to animal movement. Ecography, 40, 1369–1380. 10.1111/ecog.02801

Borchers DL, Efford M (2008) Spatially explicit maximum likelihood methods for capture–recapture studies. Biometrics, 64, 377–385. 10.1111/j.1541-0420.2007.00927.x

Chambert T, Pardo D, Choquet R, Staszewski V, McCoy KD, Tveraa T, Boulinier T (2012) Heterogeneity in detection probability along the breeding season in Black-legged Kittiwakes: implications for sampling design. Journal of Ornithology, 152, 371–380. 10.1007/s10336-010-0542-8

de Valpine P, Turek D, Paciorek CJ, Anderson-Bergman C, Lang D-C, Bodik R (2017) Programming with models: writing statistical algorithms for general model structures with NIMBLE. Journal of Computational and Graphical Statistics, 26, 403–413. 10.1080/10618600.2016.1172487

Dey S, Bischof R, Dupont PP, Milleret C (2022) Does the punishment fit the crime? Consequences and diagnosis of misspecified detection functions in Bayesian spatial capture–recapture modeling. Ecology and Evolution, 12, e8600. 10.1002/ece3.8600

Efford M (2004) Density estimation in live-trapping studies. Oikos, 106, 598–610. 10.1111/j.0030-1299.2004.13043.x

Efford MG (2022) secr: Spatially explicit capture-recapture models. R package version 4.5.6. https://CRAN.R-project.org/package=secr

Efford MG, Borchers DL, Mowat G (2013) Varying effort in capture–recapture studies. Methods in Ecology and Evolution, 4, 629–36. 10.1111/2041-210X.12049

Fiske I, Chandler R (2011) Unmarked: an R package for fitting hierarchical models of wildlife occurrence and abundance. Journal of Statistical Software, 43, 1–23. 10.18637/jss.v043.i10

Flanders NP, Gardner B, Winiarski KJ, Paton PW, Allison T, Connell AF (2015) Key seabird areas in southern New England identified using a community occupancy model. Marine Ecology Progress Series, 533, 277–290. 10.3354/meps11316

Furnas BJ, McGrann MC (2018) Using occupancy modeling to monitor dates of peak vocal activity for passerines in California. The Condor, 120, 188–200. 10.1650/CONDOR-17-165.1

Gimenez O, Viallefont A, Charmantier A, Pradel R, Cam E, Brown CR, Anderson MD, Brown MB, Covas R, Gaillard J-M (2008) The risk of flawed inference in evolutionary studies when detectability is less than one. The American Naturalist, 172, 441–448. 10.1086/589520

Kervellec M, Milleret C, Vanpé C, Quenette P-Y, Sentilles J, Palazón S, Jordana IA, Jato R, Irurtia MME, Gimenez O (2023) Integrating opportunistic and structured non-invasive surveys with spatial capturerecapture models to map connectivity of the Pyrenean brown bear population. Biological Conservation, 278, 109875. 10.1016/j.biocon.2022.109875

Kéry M, Royle JA (2009) Modeling Demographic Processes in Marked Populations. In: Inference about species richness and community structure using species-specific occupancy models in the national Swiss breeding bird survey MHB Environmental and Ecological Statistics, vol 3. (eds Thomson DL, Cooch EG, Conroy MJ), pp. 639–656. Springer, Boston, MA, USA.

Kéry M, Royle JA (2015) Applied hierarchical modelling in ecology—Modeling distribution, abundance and species richness using R and BUGS. Academic Press, New York, NY, USA.

Kéry M, Royle JA, Meredith M (2022) AHMbook: Functions and Data for the Book “Applied Hierarchical Modeling in Ecology” Vols 1 and 2. R package version 0.2.6. https://CRAN.R-project.org/package=AHMbook

Lincoln FC (1930) Calculating waterfowl abundance on the basis of banding returns. United States Department of Agriculture, Washintong D.C., USA.

MacKenzie DI, Nichols JD, Lachman GB, Droege S, Andrew Royle J, Langtimm CA (2002) Estimating site occupancy rates when detection probabilities are less than one. Ecology, 83, 2248–2255. 10.1890/0012-9658(2002)083[2248:ESORWD]2.0.CO;2

MacKenzie DI, Nichols JD, Royle JA, Pollock KH, Bailey L, Hines JE (2017) Occupancy estimation and modeling: inferring patterns and dynamics of species occurrence. Academic Press, Burlington, MA, USA.

Moqanaki EM, Milleret C, Tourani M, Dupont P, Bischof R (2021) Consequences of ignoring variable and spatially autocorrelated detection probability in spatial capture-recapture. Landscape Ecology, 36, 2879–2895. 10.1007/s10980-021-01283-x

Murtaugh PA (2009) Performance of several variable-selection methods applied to real ecological data. Ecology Letters, 12, 1061–1068. 10.1111/j.1461-0248.2009.01361.x

Nichols JD, Karanth KU (2002) Statistical concepts: assessing spatial distributions. In: Monitoring tigers and their prey: A manual for wildlife researchers, managers and conservationists in tropical Asia (eds Karanth KU, Nichols JD), pp. 29–38. Centre for Wildlife Studies, Bangalore, India.

Otis DL, Burnham KP, White GC, Anderson DR (1978) Statistical inference from capture data on closed animal populations. Wildlife Monographs, 62, 3–135.

Pollock KH, Nichols JD, Brownie C, Hines JE (1990) Statistical inference for capture-recapture experiments. Wildlife Monographs, 107, 3–97.

Pollock KH, Nichols JD, Simons TR, Farnsworth GL, Bailey LL, Sauer JR (2002) Large scale wildlife monitoring studies: statistical methods for design and analysis. Environmetrics, 13, 105–119. 10.1002/env.514

R Core Team (2022) R: A language and environment for statistical computing. R Foundation for Statistical Computing, Vienna, Austria.

Richmond OMW, Hines JE, Beissinger SR (2010) Two-species occupancy models: a new parameterization applied to co-occurrence of secretive rails. Ecological Applications, 20, 2036–2046. 10.1890/09-0470.1

Riddle JD, Mordecai RS, Pollock KH, Simons TR (2010) Effects of Prior Detections on Estimates of Detection Probability, Abundance, and Occupancy. The Auk, 127, 94–99. 10.1525/auk.2009.09062

Royle JA, Chandler RB, Sollmann R, Gardner B (2014) Spatial capture-recapture. Academic Press, Waltham, MA, USA.

Royle JA, Dorazio RM (2008) Hierarchical modeling and inference in ecology: the analysis of data from populations, metapopulations and communities. Academic Press, London, UK.

Royle JA, Fuller AK, Sutherland C (2016) Spatial capture–recapture models allowing Markovian transience or dispersal. Population ecology, 58, 53–62. 10.1007/s10144-015-0524-z

Royle JA, Nichols JD (2003) Estimating abundance from repeated presence–absence data or point counts. Ecology, 84, 777–790. 10.1890/0012-9658(2003)084[0777:EAFRPA]2.0.CO;2

Royle JA, Young KV (2008) A hierarchical model for spatial capture–recapture data. Ecology, 89, 2281–2289.

Schmid H, Zbinden N, Keller V (2004) Überwachung der Bestandsentwicklung häufiger Brutvögel in der Schweiz. Schweizerische Vogelwarte, Sempach, Switzerland.

Sollmann R (2023a) Mt or not Mt: Temporal variation in detection probability in spatial capture-recapture and occupancy models; Supplementary Information. Zenodo [Other], 10.5281/zenodo.8221229

Sollmann R (2023b) rasrage/Sollmann_2023_PCJ: Revisions (v1.1). Zenodo [Software], 10.5281/zenodo.10053390

Sollmann R, Furtado MM, Gardner B, Hofer H, Jácomo AT, Tôrres NM, Silveira L (2011) Improving density estimates for elusive carnivores: accounting for sex-specific detection and movements using spatial capture–recapture models for jaguars in central Brazil. Biological Conservation, 144, 1017–1024. 10.1016/j.biocon.2010.12.011

Strebel N, Kéry M, Schaub M, Schmid H (2014) Studying phenology by flexible modelling of seasonal detectability peaks. Methods in Ecology and Evolution, 5, 483–490. 10.1111/2041-210X.12175

Turek D, Milleret C, Ergon T, Brøseth H, Dupont P, Bischof R, De Valpine P (2021) Efficient estimation of largescale spatial capture–recapture models. Ecosphere, 12, e03385. 10.1002/ecs2.3385

Tyre AJ, Tenhumberg B, Field SA, Niejalke D, Parris K, Possingham HP (2003) Improving precision and reducing bias in biological surveys: estimating false-negative error rates. Ecological Applications, 13, 1790–1801. 10.1890/02-5078

Wegan MT, Curtis PD, Rainbolt RE, Gardner B (2012) Temporal sampling frame selection in DNA-based capture–mark–recapture investigations. Ursus, 23, 42–51. 10.2192/URSUS-D-11-00013.1

Wiest WA, Shriver WG (2016) Survey frequency and timing affect occupancy and abundance estimates for salt marsh birds. The Journal of Wildlife Management, 80, 48–56. 10.1002/jwmg.963

Williams BK, Nichols JD, Conroy MJ (2002) Analysis and management of animal populations. Academic Press, San Diego, CA, USA.

